## Supplemental legends for "Fine-tuning spatial-temporal dynamics and surface receptor expression support plasma cell-intrinsic longevity"

Supplemental Figure 1. Development of Blimp1-ERT2-Cre-TdTomato (BEC) mice and validation of the fidelity of fluorescent reporter. Related to Figure 1.

(A), Schematic of the CRISPR-Cas9 knock-in strategy to create the BEC allele, with approximate positions of and single guide RNA and genotyping primers shown. Examples of 2% agarose gel electrophoresis showing the PCR product at 277 bp in size using the genotyping primers amplified in the mutant allele compared to that in the wildtype allele. (B), FACS gating strategy showing the high fidelity of fluorescent TdTomato reporter under the endogenous *prdm1* promoter.

Supplemental Figure 2. LLPC enrichment by a multiplexed antibody panel. Related to Figure 3.

(A-C), The fold change of the expression level (gMFI) of surface markers (CD93 (A), CD81 (B), CD138 (C)) on TdTomato+YFP+ LLPCs relative to TdTomato+YFP- bulk PCs in individual mouse at indicated timepoints post tamoxifen treatment in the bone marrow (upper panel) and spleen (lower panel) of young and middle-aged mice. (D), FACS gating strategy for LLPC enrichment using a combination of 1-6 markers (one example per each number of markers used) in the bone marrow of BEC-YFP mouse at day 90 post tamoxifen treatment. Numbers represents the percentage of YFP^+^ PCs in total PCs. All bars show mean ± SEM (A-C). *, P< 0.05; **, P< 0.01; ***, P< 0.001; ****, P< 0.0001; ns, nonsignificant by unpaired Student’s t test. All graphs show pooled data from at least two independent experiments. (A-C, n = 3-7)

Supplemental Figure 3. Transmission electron microscopy of LLPCs. Related to Figure 3.

(A), Examples of transmission electron microscopy images in LLPCs and bulk PCs at day 90 post tamoxifen treatment. Scale bar is 2 μm. (B-H), Quantification of cell size (B), cytoplasm size (C), total mitochondrial area (D), normalized mitochondrial area (E), nucleus size (F), compact chromatin size (G), and normalized compact chromatin (H) in LLPCs and bulk PCs in the bone marrow and spleen. All bars show mean ± SEM. *, P< 0.05 by unpaired unpaired Student’s t test. All data are pooled data from two to three independent experiments. (B, n = 36-93; C, n = 36-99; D-E, n = 36-91; F, n = 23-77; G, n =24-69; H, n = 24-75)

Supplemental Figure 4. CXCR4 controls durable humoral response by promoting PC survival and retention in the BM. Related to Figure 4.

(A), Overlayed histograms (left panel) of the distribution of CXCR4 expression level in bulk PCs and LLPCs in the bone marrow and spleen at day 60 post tamoxifen treatment, which is quantified on the right panel. (B), FACS pseudo color plot showing the percentage of NP-specific PCs in LLPCs or bulk PCs in the bone marrow and spleen, which is quantified in (C). (D), FACS plot showing the gating strategy for each PC compartment and the percentage of labeled PCs at indicated timepoints post tamoxifen treatment in the spleen (upper panel) and bone marrow (lower panel), which is quantified in (E). (F), PC competitive competency at indicated timepoints determined by normalizing the CXCR4^cKO^:WT ratio in the bone marrow total PC compartment to that of total splenic B cell compartment (left panel) or the CXCR4^cKO^:WT ratio in the spleen total PC compartment to that of total splenic B cell compartment (right panel). All bars show mean ± SEM. *, P< 0.05; **, P< 0.01; ****, P< 0.0001; ns, non-significant by unpaired Student’s t test (A, C) or one-way ANOVA with multiple comparison correction using the Holm-Šídák test (F). Each symbol in all plots represents one mouse. All data are pooled data from two independent experiments. (A, n = 10-11; C, n = 7; E-F, n = 8-10)

Supplemental Figure 5. Analysis of public clones in PC samples. Related to Figure 7. (A). Analysis of inter-mouse clonal overlap frequencies (%), comparing each sample from young mice (i) to all other samples within groups from the remaining samples (ii). X axis labels are formatted as such (i, ii). (B). Top public clones are analyzed for their abundance (0-1, where 1 indicates found in all samples) in all PC samples (black bars), LLPC samples (red bars), and in blue, LLPC enrichment (ratio of LLPC abundance/PC abundance). Arrows indicate clones with over 75% enrichment (above red dotted line), and black dotted line indicates over-represented in LLPC samples.

**SUPPLEMENTAL VIDEO LEGENDS**

Video S1 related to Figure 2C. Maturation-dependent LLPC motility by two-photon intravital imaging in the bone marrow. Left side, intra-vital two-photon 3D time-lapse of plasma cells in the tibia, of Blimp1-YFP BEC Rosa26TdTomato mice, on day 60 post TAM. YFP+ bulk PCs (green), Tomatobright YFP+ LLPCs (yellow). In red are non-specific TdTomato expression from the Rosa26 promoter. Right side, raw movie processed and analysis of bulk PCs (green) and LLPCs (yellow) cell tracks.
