## Supplementary figures and images for "Fine-tuning spatial-temporal dynamics and surface receptor expression support plasma cell-intrinsic longevity"

### Figures S1-S5

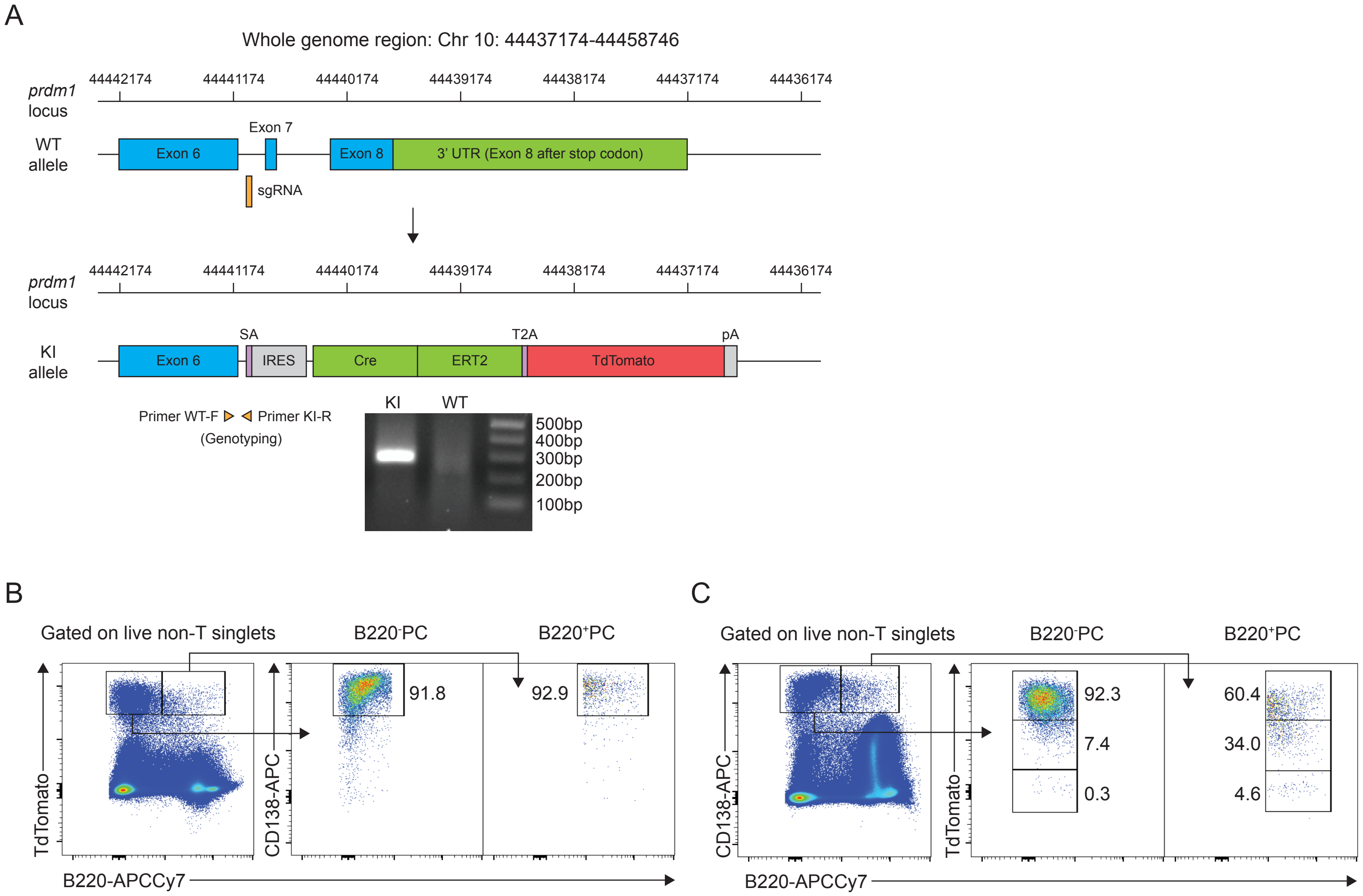

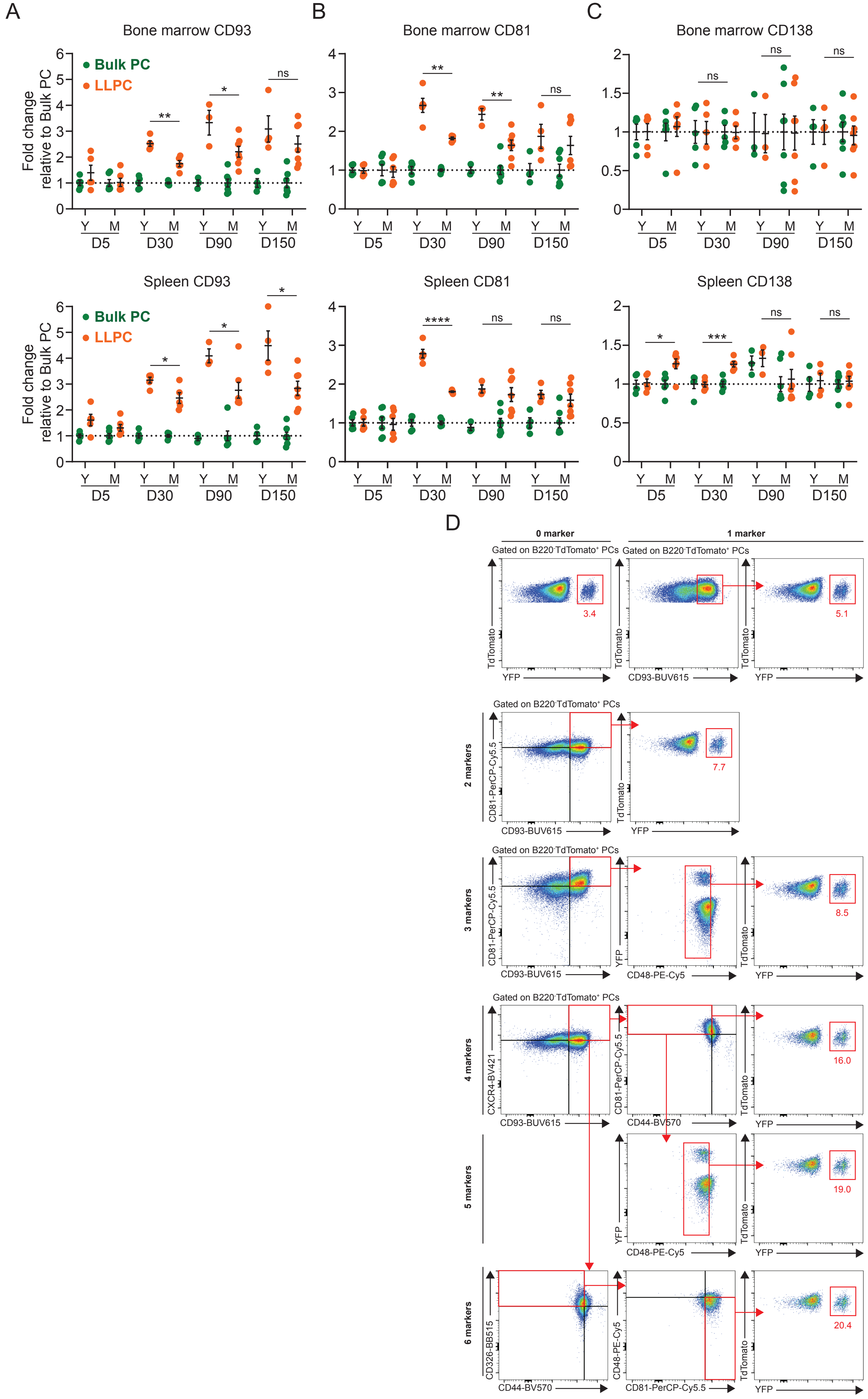

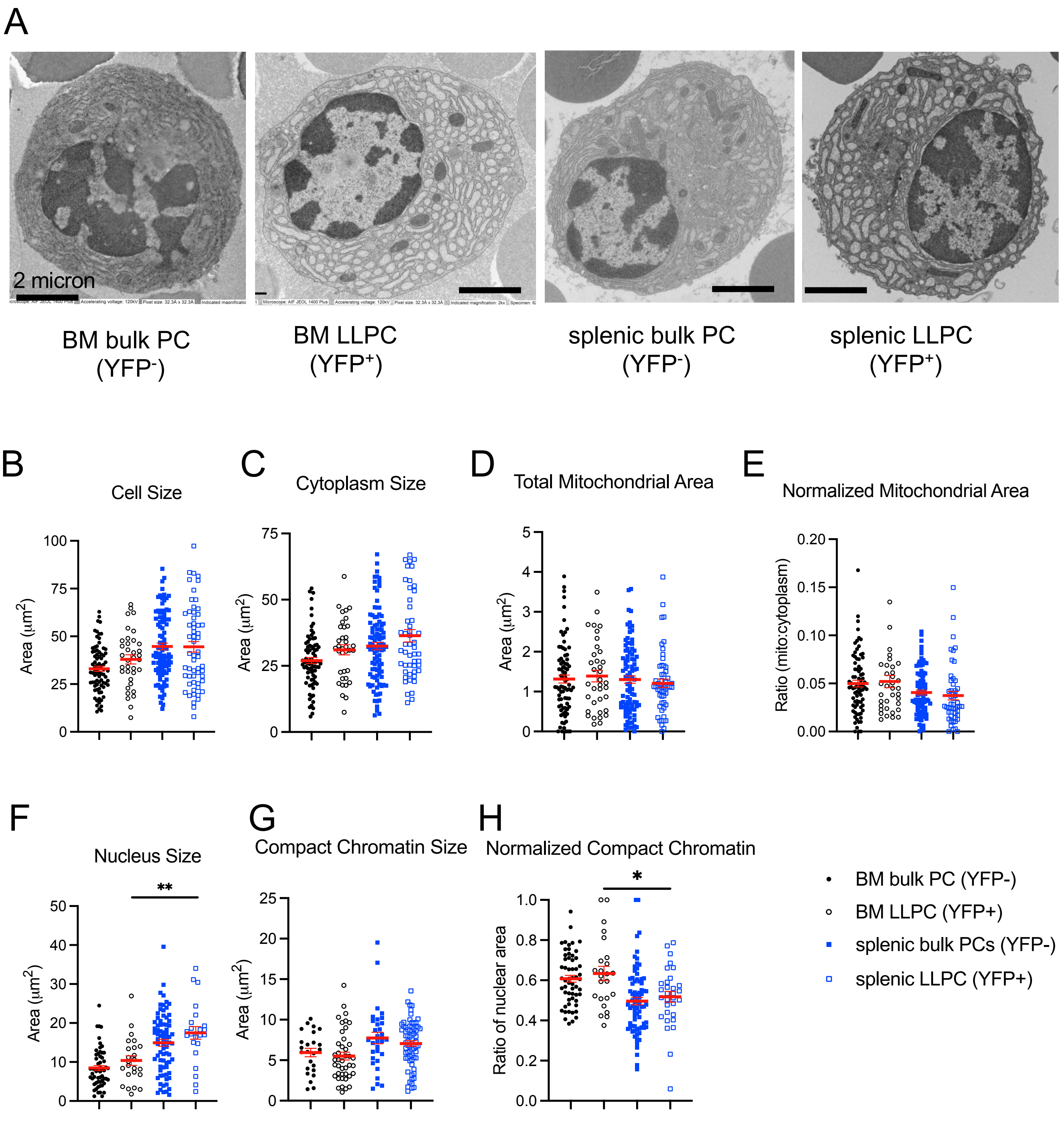

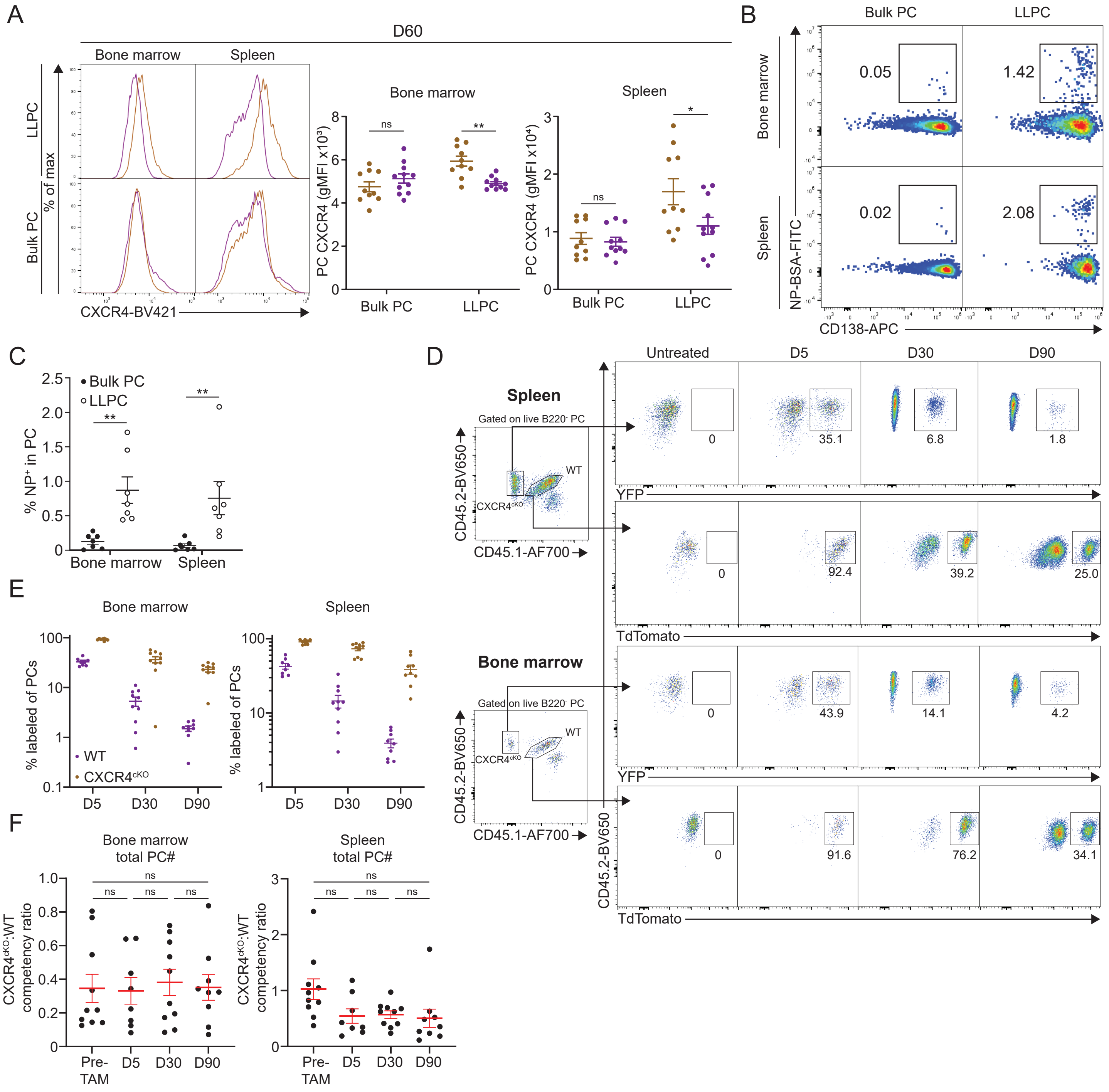

Supplemental Figure 5

A

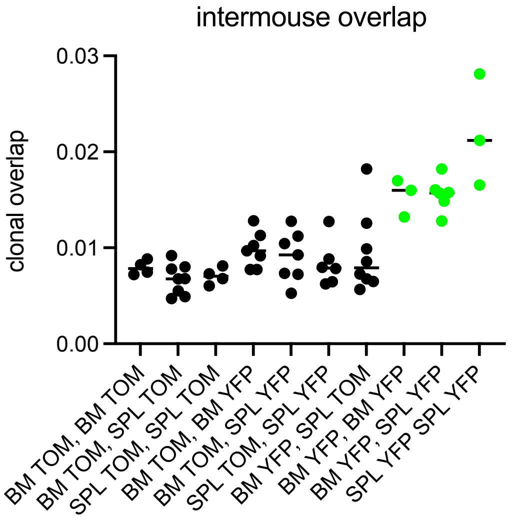


young mice only

B

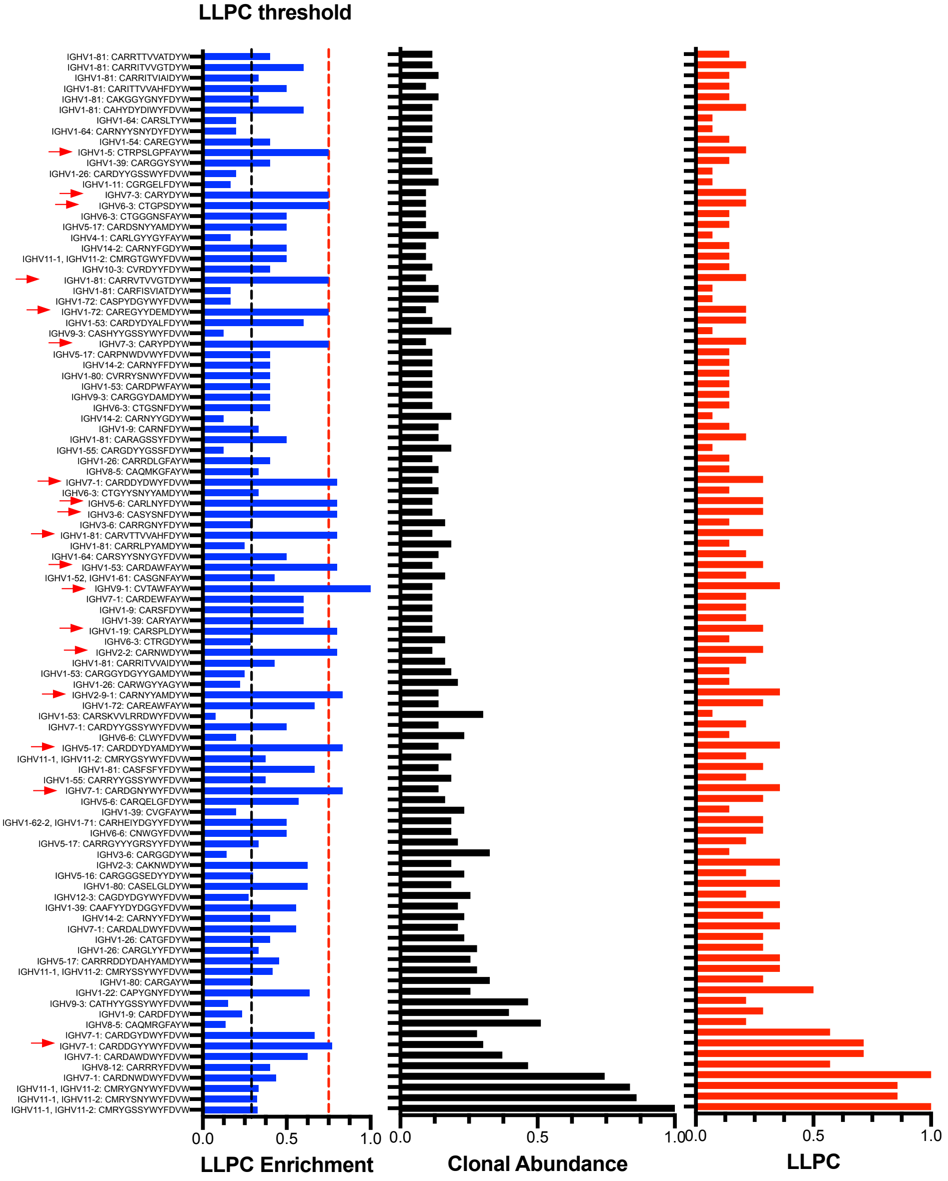
